# Feeling the beat: Temporal and spatial perception of heartbeat sensations

**DOI:** 10.1101/2020.07.27.222810

**Authors:** Sophie Betka, Marta Łukowska, Marta Silva, Joshua King, Sarah Garfinkel, Hugo Critchley

**Author notes:** Corresponding author: Dr Sophie Betka. Corresponding Author @DoctoresseSoso.

## Abstract

The majority of interoceptive tasks (i.e. measuring the sensitivity to bodily signals) are based upon the heartbeat sensations perception. However, temporal perception of heartbeats varies among individuals and confidence and spatial perception of heartbeats in relation to temporal judgements have not yet been systematically investigated. This study examined the perception of heartbeat sensations in both time and spatial domains, using a multi-interval heartbeat discrimination task. The relationship between these domains was investigated, as well as the contribution of mental health conditions and cardiovascular parameters. Heartbeat sensations occurred on average ~250ms after the ECG R-wave and were more frequently sampled from the left part of the chest. Participants’ confidence in their experience of heartbeat sensations was maximal for the 0 ms interval. Interestingly, higher confidence was related to reduced dispersion of sampling locations but we found evidence toward the absence of relationship between temporal and spatial heartbeat sensations perception, using Bayesian statistics. Finally, we found evidence toward a relationship between spatial precision of heartbeat sensations and state anxiety score, which seems independent from the cardiovascular parameters. This systematic investigation of heartbeat sensations perception provides important fresh insights, informing the mechanistic understanding of the interoceptive signaling contribution to emotion, cognition and behaviour.

## Introduction

Interoception is the sense of the internal state of the body, and includes the perception of bodily signals coming from the viscera or glands (Cameron 2001; Sherrington 1907). Interoception conveys and represents essential physiological information concerning health (including internal sensations of somatic pain and bodily temperature), and gives rise to motivational feelings including experience of thirst and hunger. Neurally, afferent signals travel via vagus nerve and spinal (e.g. Lamina I spinothalamocortical) pathways to brainstem and thalamus, then relayed notably to insular cortex (Craig 2002). A further projection into anterior insula gives rise to an integrated representation of these bodily signals that is accessible to conscious appraisal and a dynamic substrate for subjective feelings (Critchley 2004).

Interoceptive signals for the most part inform automatic, unconscious homeostatic reflexes (Jänig 2006). Moreover, people vary in their capacity for conscious access to interoceptive sensations, and these individual differences are considered relevant to the experience of emotions and vulnerability to pathological psychological and somatic symptoms (Quadt, Critchley, and Garfinkel 2018). Consequently, research has focused on the objective measurement of interoception, using behavioral tasks through which interoceptive accuracy can be quantified from performance (Garfinkel et al. 2015). Most widely, such tasks use the perception cardiac sensations at rest: heartbeats are clear and discrete events, which are easily measurable (Betka et al., 2018) and relevant ‒the changing strength and timing of heartbeats are signatures of changing physiological arousal that accompany relevant emotions, exercise, injury, or illness. Hence, tests of heartbeat perception tasks dominate, providing a baseline metric of interoceptive sensitivity. There has been less interest in the source of the heartbeat sensation, which accompanies the ejection of blood from the heart into the aorta at ventricular systole, and includes physical changes within the vessels, chest and body (including the somatosensory, quasi-interoceptive, hitting of the inner chest wall by the heart). Nevertheless, some cardiac interoception tasks rely on the participant discriminating the timing of own heartbeats relative to a phasic external stimulus, such as an auditory tone or flashing light. This approach raises possible confounds, since different people might perceive heartbeats through different sensory channels and hence show variability in when and where they perceive their own heartbeats (Brener and Kluvitse 1988; Brener, Liu, and Ring 1993; Brener and Ring 2016; Ring and Brener 1992; Wiens and Palmer 2001). One channel of interoceptive cardiac information comes from the phasic firing of specialized arterial (aortic and carotid) baroreceptors, as blood ejected from the heart stretches the vessel walls (Garfinkel and Critchley 2016). Some earlier researchers estimated, across individuals, the average delay between the ventricular contraction, peak baroreceptors activation and the heartbeat sensation to be approximately 150ms (Whitehead et al. 1977). More systematic investigations of the temporal perception of heartbeat sensations indicate that people judge external auditory tones to be most simultaneous with heartbeat sensations if presented between 100 ms and 300 ms after the electrocardiogram (ECG) R-wave, the signature of myocardial electrical depolarization triggering ventricular contraction. Typically, the mean or the median of chosen temporal intervals is qualified as the temporal location of heartbeat sensation which lies between 228 and 288 ms after the R-wave (Brener et al. 1993; Brener, Ring, and Liu 1994; Ring and Brener 1992; Schneider, Ring, and Katkin 1998; Wiens and Palmer 2001).

However, such results do not only represent ‘pure’ interoceptive information conveyed to the brain via the vagal and spinal afferent pathways. Indeed, despite full denervation of the heart and aortic arch, some heart transplant recipients may accurately feel their heartbeats (Barsky et al. 1998). Similarly, on a heartbeat detection task, a patient with both an extracorporeal left ventricular assist device and an endogenous heart was rather following artificial pump-beats (via abdomen somatosensory feedback) than his actual endogenous heartbeats (Couto et al. 2014). Finally, a patient with extensive bilateral damage to insula and anterior cingulate cortex -structures underlying interoceptive processes (Critchley et al. 2004)- showed preserved interoceptive accuracy after the bolus administration of isoproterenol (Sahib S. Khalsa et al. 2009). Only after anaesthetising the patient’s chest in the region of maximal heartbeat sensation, interoceptive awareness was impaired (Sahib S. Khalsa et al. 2009). Together, such results suggest that the somatosensory pathway also contributes to heartbeat sensations.

The spatial location of heartbeat sensations has not been studied as extensively as the temporal aspect. Khalsa and colleagues asked participants to trace on a manikin template (representing their own body) the location of their heartbeat sensations. During low arousal states, participants mostly felt their heartbeats in the lower left chest. Some participants also reported heartbeat sensations in the head, neck, belly and arms (Hassanpour et al. 2016; Khalsa et al. 2018; Khalsa et al. 2009; Khalsa et al. 2009). Good heartbeat perceivers, based on how accurately they performed the heartbeat detection ‘counting’ task (in which the reported number of ‘felt’ heartbeats, counted over different time periods are compared to veridical heartbeats, measured using ECG) report more spontaneous sensations (SPS; e.g. tickling, tingling or even warming sensations) in the hands than poor heartbeat perceivers (Michael et al. 2015). The time interval from the ECG R wave to the finger pulse (pulse transit time) is typically estimated to ~250ms (Ma and Zhang 2005). However, microneurography reveals firing of mechanoreceptors in the finger’s to occur as early as 200ms after ECG R-wave (Macefield 2003). Nevertheless, the relationship between temporal and spatial locations of heartbeat sensations has not yet been explored.

Interestingly, interoceptive abilities are likely impaired across different mental health conditions. It is likely that both temporal and spatial locations of heartbeat sensations may be affected in these groups (Betka et al. 2017, 2018; Khalsa et al. 2018; Paulus and Stein 2010; Quadt et al. 2018). Similarly, individual differences in physical constitution, notably body mass index (BMI) (Brener and Kluvitse 1988; Brener et al. 1993; Michael et al. 2015; Ring and Brener 1992, 2018; Rouse, Jones, and Jones 1988; Wiens and Palmer 2001) and cardiovascular parameters (e.g. interbeat intervals, heart rate, heart rate variability) (Brener et al. 1993; Knapp-Kline and Kline 2005; Ring and Brener 1992) may shape when and where heartbeat sensations are perceived.

## Aims

The present study aimed to examine in detail the perception of heartbeat sensations, using a multi-interval heartbeat discrimination task (Brener and Kluvitse 1988), to address the following questions:

- *When* do people perceive their own heartbeats with confidence? We quantified the timing of heartbeat perception using a simultaneity judgement of heartbeat sensation relative to the presentation of an external auditory tone, triggered at different delays (SOAs) from the ECG R-wave. We predicted that participants would judge tones delivered between 100-300ms after R-wave as simultaneous with their heartbeat (Brener and Kluvitse 1988; Brener et al. 1993, 1994; Ring and Brener 1992).
- *Where* do people feel their heartbeat with confidence? On a two-dimensional body map, participants marked the anatomical location of maximal heartbeat sensation. We predicted that participants would typically feel their heartbeat in the left chest and neck areas (Khalsa et al. 2018; S. S. Khalsa et al. 2009).
- Does the timing of heartbeat perception relate to the spatial location of heartbeat perception? We tested if the observed timing of heartbeat sensation was predicted by the anatomical location and spatial dispersion of the heartbeat sensation.
- What influences individual differences in the temporal and spatial perception of heartbeats? We tested for relationships between temporal and spatial differences in heartbeat perception and somatic (BMI), affective (Anxiety, Depression, Alexithymia), and physiological measures (inter-beat interval, heart rate variability). We hypothesized that BMI impact neither temporal nor spatial aspect of heartbeat sensations (Brener and Kluvitse 1988; Brener et al. 1993; Ring and Brener 1992, 2018). However, we predict that individual differences in affective functioning (Anxiety, Depression, Alexithymia) will influence both aspects of heartbeat sensation (Betka et al. 2017, 2018; Khalsa et al. 2018; Paulus and Stein 2010). Finally, we expect that heart rate will impact heartbeat perception ‒namely, slower heart rate (i.e. longer inter-beat intervals) will be associated with better heartbeat perception (Knapp-Kline and Kline 2005).

## Results

Here, we examined the perception of heartbeat sensations, using a multi-interval heartbeat discrimination task (J. Brener et al., 1993; see Figure 1 and the methods section). On each trial, the participant listened to a sequence of 5 tones, in which were all presented either 0, 100, 200, 300, 400 or 500 ms after the R-wave on ECG. After listening to an individual sequence, the participant decided whether or not the tones were played simultaneously with perceived own heartbeats. The participant then rated confidence in that decision, using a visual analogue scale (from 0=not at all to 100=completely). The participant was then presented with an outline image of a body on the screen and where asked to mark on the body template where did they feel their heartbeat sensation the strongest. Descriptive and Bayesian statistics alongside traditional mixed-effects linear models, contrasts and correlations were used to answer the following questions.

- ***When do people perceive their heartbeats in relation to an external auditory tone and how confident are they?***
- ***The temporal locations of heartbeat sensations happen around 250ms after the ECG R-wave*.**

**Figure 1.**
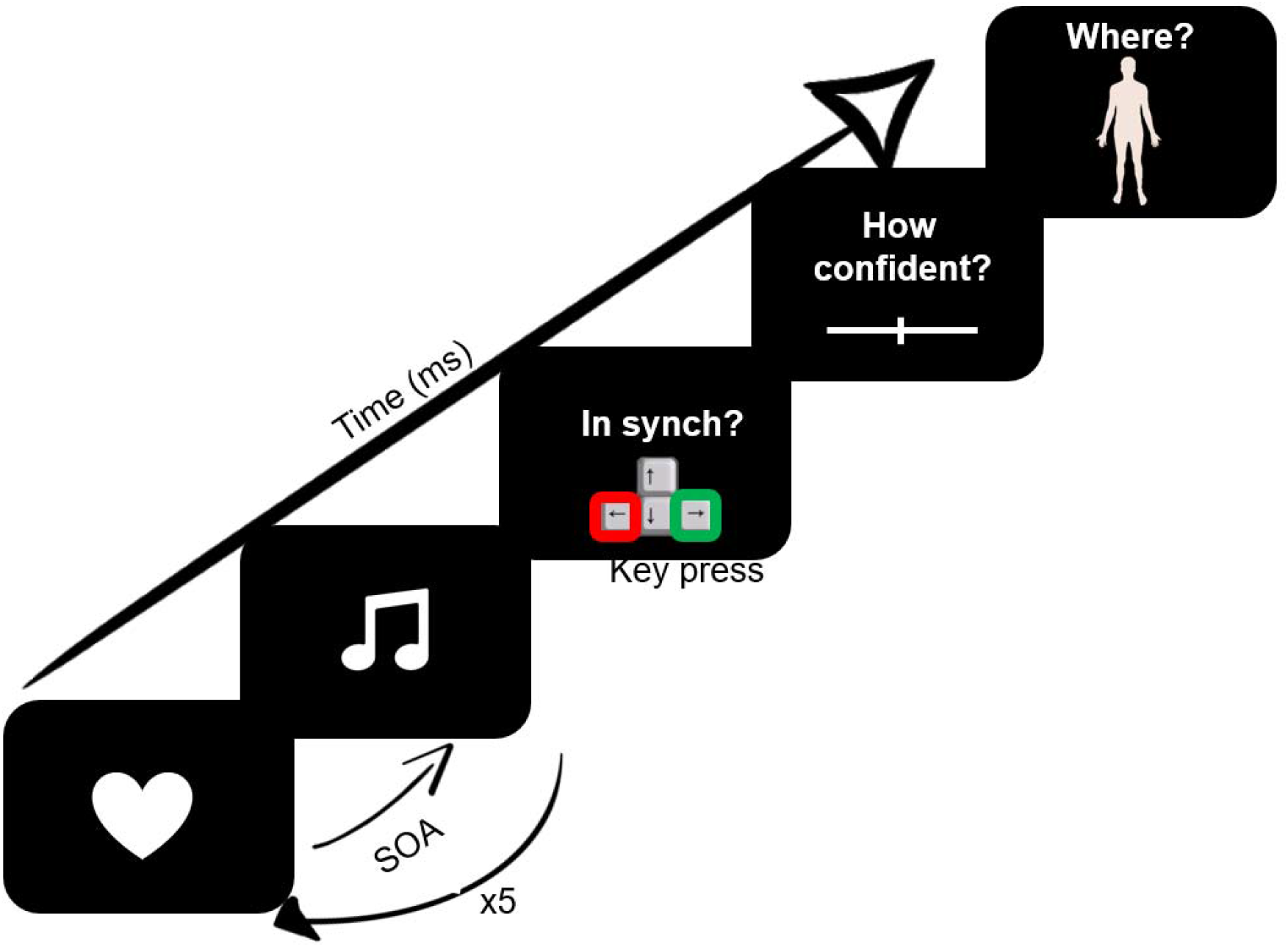
Schematic representation of a trial of the multi-interval heartbeat discrimination task. On each trial, the participant’s ECG R-peak was the priming trigger for tone presentations at a specific delay (0 to 500ms), repeated 5 times per trial. The participant then judged the perceived simultaneity of the tones with their own heartbeats, rated confidence in that judgement, and marked where on the body the heartbeat sensation was felt.

On average, participants felt their heartbeat 257.40ms ±31.31ms (*Median* = 258.70 ± 55.77) after the actual ECG R-peak (see Table S1). Their modal preferred interval was 300ms and ranged from 0 to 500ms; its mean was equal to 265.40ms. The specificity of discrimination (standard deviation of the modal preferred interval) (Brener and Ring 2016) was 149.36ms. Individual performance is depicted on Figure S1. These findings confirm predictions generated by previous work (Brener et al. 1993, 1994; Ring and Brener 1992; Wiens and Palmer 2001)

**Figure 2.**
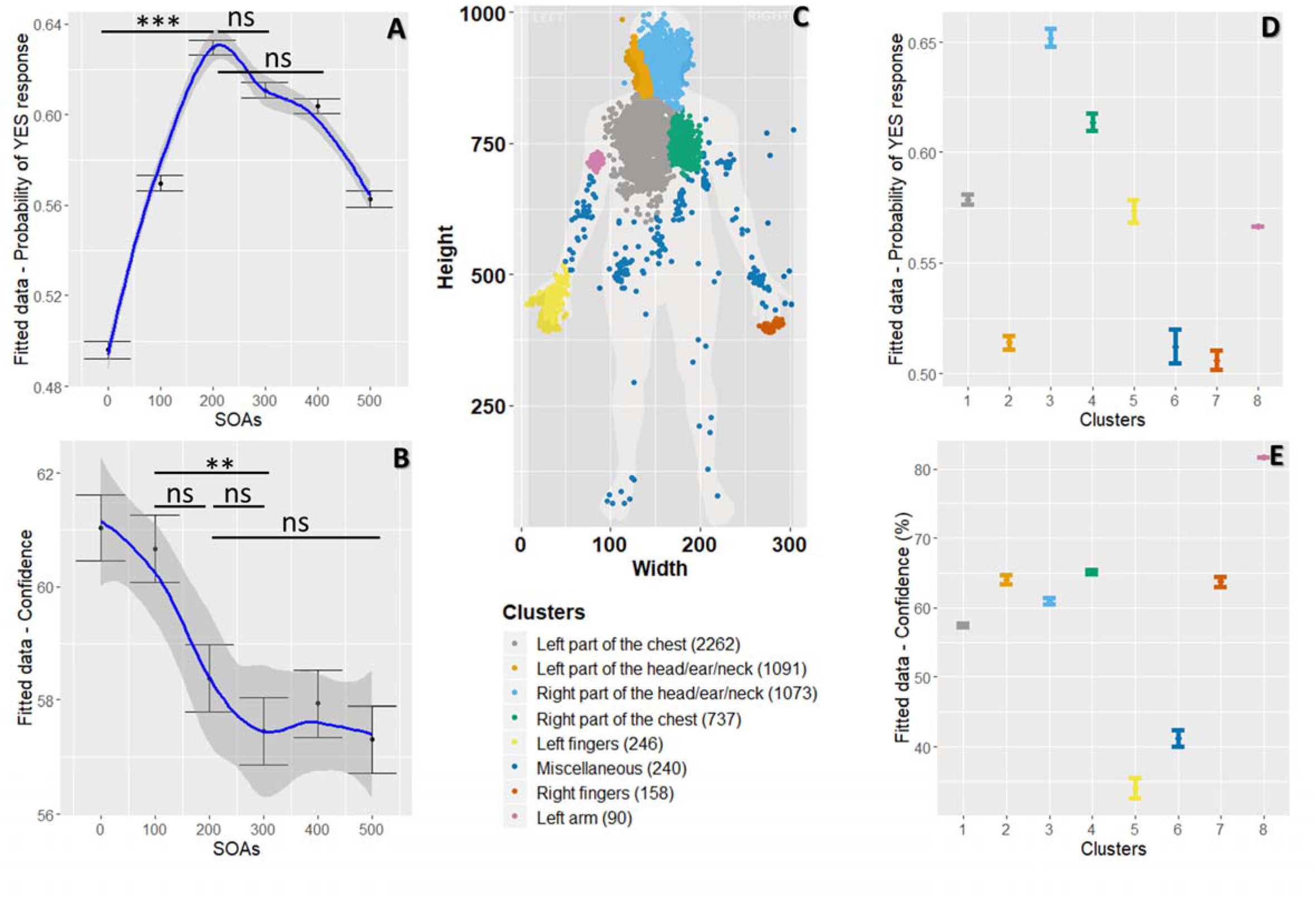
Results from the temporal and spatial heartbeat perception. **A** Temporal perception of heartbeat sensations: Graph of the effect of delays (stimulus onset asynchrony; SOAs) on the probability of yes response to simultaneity judgments of heartbeat with auditory tone (showing mean and error bars representing standard errors), while taking in account within-participant variability. **B** Confidence in the temporal perception of heartbeat sensations: Graph of the effects of delays (SOAs) on confidence ratings (with mean and error bars representing standard errors), while considering within-participants variability. **C** Visual representation of the eight clusters where participants reported sampling their heartbeat sensations. Each dot represents a trial. (1 = Left part of the chest, 2 = Left part of the head/ear/neck, 3 = Right part of the head/ear/neck, 4 = Right part of the chest, 5 = Left fingers, 6 = Miscellaneous, 7 = Right fingers, and 8 = Left arm).The number of observations is written in brackets for each cluster. **D**. Graph of the effect of clusters on the probability of yes response (with mean and error bars representing standard errors), while considering within-participants variability. **E**. Graph of the effect of clusters on confidence ratings (with mean and error bars representing standard errors), while considering within-participant variability. Significance of the main results are indicated.by ns = non-significant; \*\**p-*values < .01; \*\*\**p-* values < .001.

Results of the mixed-effects regression model provide further insight (see Figure 2.A and Table S2) We report 95% confidence interval (95% CI), Savage-Dickey density ratio Bayes Factor (*BF*), the most credible value, and 95% Credible Interval (95% CrI). A delay of 200ms from R-wave produced the highest probability of answering “Yes” for a judgment of simultaneity (β = 0.59, *SE* = 0.11, 95% CI [0.40, 0.76], *p*-value < .001, *BF* = 252.44, 62.84%, 95% CrI [58.78, 66.99]), compared to the non-delay condition (β = −0.007, *SE* = 0.141, 95% CI [−0.20, 0.20], *p*-value = .961, *BF* = 0.01, 49.66%, 95% CrI [45.62, 53.90]).

Based on previous work, we predicted that participants would judge tones delivered between 100-300ms after R-wave as simultaneous with their heartbeat (Brener and Kluvitse 1988; Brener et al. 1993, 1994; Ring and Brener 1992). Planned contrasts between delays from 100 to 300 ms were therefore computed. Interestingly, no difference in terms of probability of answering “Yes” was observed for a delay of 100ms compared to 200ms (β = −0.27, *SE* = 0.09, 95% CI [−0.50, −0.03], corrected *p*-value <. 05, *BF* = 0.3, 6.05%, 95% CrI [0.27, 11.84]), and compared to the 300ms delay condition (β = −0.18, *SE* = 0.09, 95% CI [−0.41, 0.05], corrected *p*-value = .157, *BF* = 0.04, −4.25%, 95% CrI [−9.84, 1.90]). No difference between the 200ms and the 300ms delay conditions was observed (β = 0.09, *SE* = 0.09, 95% CI [−0.15, 0.32], corrected *p*-value = .418, *BF* < 0.01, 1.89%, 95% CrI [−3.99, 7.49]). A post-hoc contrast showed no significant difference in the probability of answering “Yes” between the 200ms and the 400ms delay conditions (β = 0.12, *SE* = 0.093, 95% CI [−0.06, 0.29], corrected *p*-value = .418, *BF* = 0.01, 2.58%, 95% CrI [−3.20, 8.47]). Our results demonstrate that participants felt their heartbeats between 100 and 400ms (*Mean* = 257.40ms); reinforcing extant literature (Ring and Brener 1992; Yates et al. 1985).

The mean of confidence ratings (on 0-100 VAS) was equal to 58.78 ± 19.32 (*Median* = 62.59, *Rang*e = 3.53 – 97.6; see Table S1). Results of the mixed-effects regression model (see Figure 2.B and Table S4) revealed that lowest confidence ratings were observed for delays of 300ms (β = −3.65, *SE* = 0.91, 95% CI [−5.43, −1.87], *p*-value = .001, *BF* > 1000, 57.47%, 95% CrI 55.67, 59.24]) and 500ms (β = − 3.75, *SE* = 0.91, 95% CI [−5.53, −1.97], *p*-value < .001, *BF* > 1000, 57.32%, 95% CrI [55.55, 59.11]), compared to the non-delay condition (β = 61.04, *SE* = 2.74, 95% CI [55.62, 66.46], *p*-value < .001, *BF* > 1000, 60.97%, 95% CrI 59.20, 62.75]).

Based on published findings we predicted that confidence in timing simultaneity would relate to perceptual ease, and therefore be maximal for the 0ms and 500ms intervals, and minimal when disCrlminating simultaneity over the intervals between 100ms to 300ms. Planned contrasts were computed: A significant difference in confidence was observed between the 100ms delay and the 300ms delay condition (β = 3.24, *SE* = 0.091, 95% CI 0.97, 5.51], corrected *p*-value < .01, *BF* = 23.25, 3.21%, 95% CrI [0.65, 5.73]). Moreover, no significant differences in confidence were observed between the 200ms delay condition the 100ms delay condition (β = 2.25, *SE* = 0.091, 95% CI [−0.03, 4.52], corrected *p*-value < .05, *BF* = 1.37, 2.26 %, 95% CrI [−0.27, 4.78]), the 300ms delay condition (β = 0.10, *SE* = 0.91, 95% CI [−1.28, 3.27], corrected *p*-value = .456, *BF* = 0.12, 0.94%, 95% CrI [−1.57, 3.48]) or the 500ms delay condition (β = −0.1, *SE* = 0.091, 95% CI −1.18, 3.37], corrected *p*-value = .456, *BF* = 0.14, 1.08%, 95% CrI [−1.45, 3.60]). Participants were thus more confident for the 0ms interval, but less for intervals between 100ms to 500ms.

- ***Where do people feel their heartbeat and how confident are they?***
- ***The spatial locations of heartbeat sensations happen in the left part of the chest*.**

After each trial, participants marked the anatomical site of maximal heartbeat sensation on a body map. The distance between the sampling location and the heart (assigned as a standardised location) was computed using coordinates marked by the participant on the body outline. Next, the mean of the distance to the heart was computed for each participant (Distance from the heart). We also computed dispersion from sampling locations by computing the mean of the standard deviation of X coordinates and the standard deviation of Y coordinates, for each participant ((sd(X) + sd(Y))/2). Finally, clusters of sampling location data points were defined using expectation-maximization algorithm for fitting mixture-of-Gaussian models (mclust R package) (Scrucca et al. 2016) and attributed to body parts and assigned names based on visual inspection. We isolated eight clusters (see Figure 2.C). The most frequently reported spatial location of heartbeat sensations (modal preferred cluster) was around the left part of the chest (cluster 1), coherent with previous work (Khalsa et al. 2018; S. S. Khalsa et al. 2009). Individual data are presented on Figure S2.

Results of the mixed-effects regression model are presented in Figure 2.D and Table S4. The highest number of simultaneity judgements were observed in fact for the right part of the head/ear/neck (β = 0.37, *SE* = 0.11, 95% CI [0.15, 0.59], *p*-value = .001, *BF* = 720.06, 65.14%, 95% CrI [61.42, 68.87]) and the right part of the chest (β = 0.48, *SE* = 0.14, 95% CI [0.21, 0.77], *p*-value = .001, *BF* = 273.05, 61.33%, 95% CrI [56.45, 66.08]) in contrast to the left part of the chest (β = 0.26, *SE* = 0.10, 95% CI [0.06, 0.46], *p*-value = .01, *BF* = 0.26, 57.82%, 95% CrI [55.08, 60.61]). Counter-intuitively, these findings indicate that, when participants sample their heartbeat sensations from the right part of their head/ear/neck or chest, a greater probability to reply “Yes” wass observed compared to when the participants sample their heartbeat sensations from their left part of the chest.

In terms of confidence, results of the mixed-effects regression model (Table Figure 2.E and Table S5) revealed that highest perceptual confidence was observed for the left part of the head/ear/neck (β = 8.15, *SE* = 1.30, 95% CI [5.6, 10.70], *p*-value < .001, *BF* > 1000, 64.06%, 95% CrI [62.34, 65.79]) compared to the left part of the chest (β = 55.75, *SE* = 2.52, 95% CI [50.78, 60.73], *p*-value < .001, *BF* > 1000, 56%, 95% CrI [56.35, 58.78]).

- ***Does the timing of heartbeat perception relate to the spatial location of heartbeat perception?***

We tested if the timing of, and confidence in, perceived heartbeat sensations was related to the distance from the heart of the indicated location of sampling, and/or the spatial dispersion of the sampling location (see Table 1). Pearson correlation coefficient, *p*-value and Bayes factor (*BF*) were computed for each relationship. Higher confidence was associated with reduced sampling location dispersion (see Figure 3.A). However, evidence toward no relationship was observed between the standard deviation of heartbeat temporal perception and distance from the heart and also between the median of heartbeat temporal perception and the sampling location dispersion. The remainder of such relationships were characterized by a *BF* between 3 and 1/3 indicating that there was insufficient evidence in either direction to make a firm conclusion (Jeffreys 1961; Lee and Wagenmakers 2013).

**Table 1.** Pearson correlation coefficients *r*, *p*-values and Bayes Factor (*BF*) for correlations between temporal (Average, standard deviation, median, mode, confidence) and spatial (Distance from the heart, sampling location dispersion) heartbeat sensation parameters. *BF* supporting evidence for a relationship (*BF* > 3) or supporting the absence of a relationship (*BF* < 1/3) between the variables are represented in bold.

|  | Distance from the heart |  |  | Sampling location dispersion |  |  |
| --- | --- | --- | --- | --- | --- | --- |
|  | *r* | *p-value* | *BF* | *r* | *p-value* | *BF* |
| **Average** | 0.151 | 0.296 | 0.521 | -0.074 | 0.611 | 0.358 |
| **Standard deviation** | -0.038 | 0.793 | **0.328** | 0.126 | 0.383 | 0.449 |
| **Median** | 0.176 | 0.222 | 0.623 | -0.033 | 0.82 | **0.326** |
| **Mode** | 0.177 | 0.218 | 0.632 | -0.047 | 0.744 | 0.334 |
| **Confidence** | -0.175 | 0.225 | 0.618 | -0.37 | 0.01 | **6.265** |

**Figure 3.**
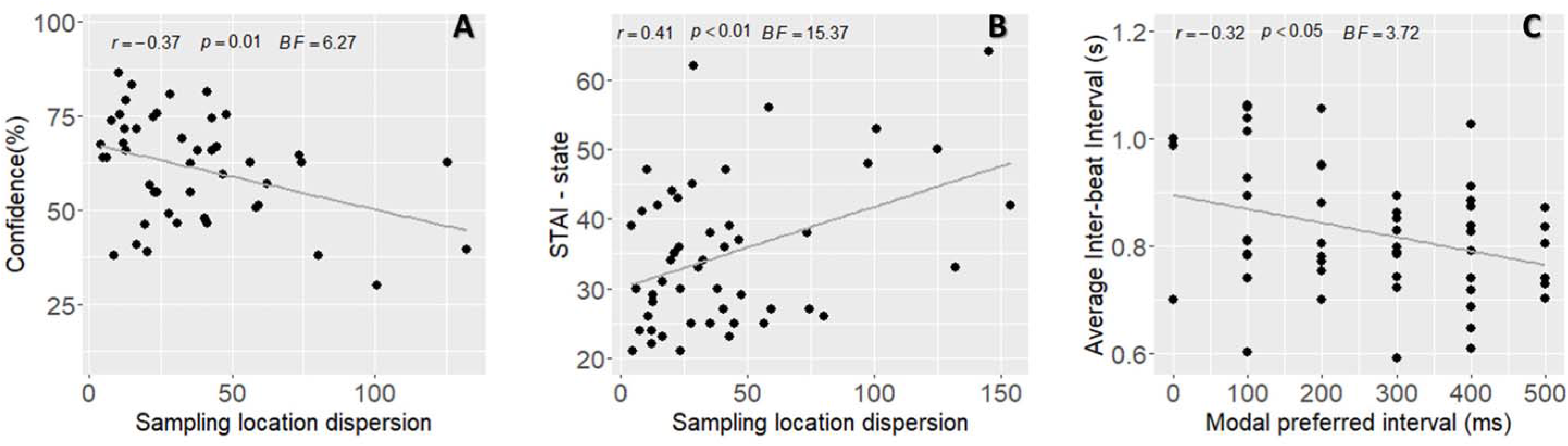
Correlations between heartbeat sensation parameters, interindividual differences and physiological measures. **A.** Pearson correlation with coefficient *r, p*-values, *BF* (Bayes Factor) for the relationship between confidence and sampling location dispersion. **B.** Pearson correlation with coefficient *r, p*-values, *BF* for the relationship between the STAI1 scores (state anxiety) and sampling location dispersion. **C.** Pearson correlation with coefficient *r, p*-values, *BF* for the relationship between the average interbeat intervals (IBI), and the modal preferred intervals.

- ***What determines individual differences in the temporal and spatial perception of heartbeats?***

Perceptual accuracy on heartbeat detection tasks has been linked to a ‘slow and steady’ heart rate (Knapp-Kline and Kline 2005) and thus diminished by increased heart rate variability (HRV and respiratory sinus arrhythmia, usually associated with slower heart rate) HRV, perhaps also through accompanying changes in stoke volume, may also decrease confidence in, and add variability to the spatial precision of heartbeat perception. In contrast, heartbeat perceptual accuracy may be increased by sampling cardiac sensations from (somatosensory) locations that can offer greater sensory precision. Finally, individual differences in body mass and emotional state may directly or indirectly influence specific aspects of heartbeat perception. Therefore, we tested for correlations between temporal and spatial differences in heartbeat perception and participants’ demographic (BMI), psychological (Anxiety, Depression, Alexithymia) and physiological measures (inter-beat interval, heart rate variability) (see Table 2 & Table 3).

**Table 2.** Pearson correlation coefficients *r*, *p*-values and Bayes Factor (*BF*) for correlations between heartbeat sensation temporal and spatial perception parameters, confidence, and psychometric/demographic parameters (BMI = body mass index; BDI = depression scores; STAI1 = anxiety state scores; STAI2 = anxiety trait scores;TAS = alexithymia scores). *BF* supporting evidence for a relationship (*BF* > 3) or supporting the absence of a relationship (*BF* < 1/3) between the variables are represented in bold.

|  | Average |  |  | Confidence |  |  | Distance from the heart |  |  | Sampling location dispersion |  |  |
| --- | --- | --- | --- | --- | --- | --- | --- | --- | --- | --- | --- | --- |
|  | *r* | *p-value* | *BF* | *r* | *p-value* | *BF* | *r* | *p-value* | *BF* | *r* | *p-value* | *BF* |
| **BMI** | -0.191 | 0.176 | 0.717 | -0.06 | 0.673 | 0.339 | 0.168 | 0.244 | 0.587 | 0.125 | 0.388 | 0.445 |
| **BDI** | -0.098 | 0.489 | 0.388 | -0.045 | 0.753 | **0.327** | 0.134 | 0.355 | 0.469 | 0.15 | 0.298 | 0.519 |
| **STAI1** | -0.059 | 0.679 | 0.338 | -0.233 | 0.097 | 1.09 | 0.285 | 0.045 | 1.942 | 0.41 | 0.004 | **15.366** |
| **STAI2** | 0.01 | 0.942 | **0.313** | -0.152 | 0.283 | 0.527 | 0.26 | 0.068 | 1.429 | 0.187 | 0.193 | 0.685 |
| **TAS** | 0.091 | 0.52 | 0.377 | 0.011 | 0.937 | **0.314** | 0.03 | 0.836 | **0.325** | 0.155 | 0.283 | 0.536 |

In this non-clinical sample, evidence supporting no relationship was observed between temporal location of heartbeat sensation and trait-anxiety scores (STAI2), between confidence, depression and alexithymia scores (BDI and TAS) and between alexithymia scores and distance from the heart. State anxiety was associated with more variable regional sampling of heartbeat sensations (see Figure 3.B). Interestingly, relationships between STAI1 and sampling location dispersion survived correction for mean inter-beat interval and HRV (Dispersion correcting for mean |B| *r* =0.394, *p* < .01; and HRV: *r* =0.385, *p* < .01). A lack of evidence in either direction to make a firm conclusion was observed for the remainder of the relationships.

**Table 3.** Pearson correlation coefficients *r*, *p*-values and Bayes Factor (*BF*) for correlations between average inter-beat interval (IBI), its variability (IBI sd, HRV), psychometric/demographic parameters (BMI = body mass index; BDI = depression scores; STAI1 = anxiety state scores; STAI2 = anxiety trait scores;.TAS = alexithymia scores)., temporal and spatial perception of heartbeat sensation and confidence. *BF* supporting evidence for a relationship (*BF* > 3) or supporting the absence of a relationship (*BF* < 1/3) between the variables are represented in bold.

|  | IBI |  |  | IBI sd (HRV) |  |  |
| --- | --- | --- | --- | --- | --- | --- |
|  | *r* | *p-value* | *BF* | *r* | *p-value* | *BF* |
| **BMI** | 0.007 | 0.963 | **0.313** | 0.13 | 0.357 | 0.46 |
| **BDI** | -0.15 | 0.289 | 0.52 | 0.052 | 0.714 | **0.332** |
| **STAI1** | -0.135 | 0.341 | 0.352 | 0.102 | 0.47 | 0.39 |
| **STAI2** | -0.073 | 0.607 | 0.471 | 0.099 | 0.484 | 0.396 |
| **TAS** | -0.175 | 0.216 | 0.626 | 0.034 | 0.812 | **0.321** |
| **Average** | -0.039 | 0.784 | **0.323** | -0.06 | 0.675 | 0.339 |
| **Standard deviation** | -0.153 | 0.28 | 0.53 | 0.046 | 0.743 | **0.328** |
| **Median** | -0.032 | 0.825 | **0.32** | -0.125 | 0.377 | 0.445 |
| **Confidence average** | 0.158 | 0.263 | 0.552 | 0.002 | 0.991 | **0.313** |
| **Mode** | -0.323 | 0.019 | **3.718** | -0.214 | 0.128 | 0.894 |
| **Distance from the heart** | 0.037 | 0.797 | **0.328** | -0.113 | 0.433 | 0.42 |
| **Sampling location dispersion** | -0.016 | 0.913 | **0.32** | 0.162 | 0.26 | 0.564 |
| **Preferred cluster** | -0.018 | 0.481 | 0.395 | 0.021 | 0.073 | 1.347 |

We also observed that participants with slower heart rate preferred shorter time intervals for temporal location of heartbeat sensation (see Figure 3.C). However, evidence supporting no relationship was observed between interbeat-interval and BMI, heartbeat perception timing, its median, distance from the heart and sampling location dispersion (see Table 3). Concerning the heart rate variability, evidence supporting the absence of a relationship was observed for depression scores, alexithymia, standard deviation of heartbeat perception timing, and confidence. A lack of evidence in either direction to make a firm conclusion was observed for the rest of the relationships.

To sum up, we found that heartbeat sensations occurred on average 250ms after the ECG R-wave and were more frequently sampled from the left part of the chest. Individuals who felt heartbeats on the right of their upper body (head/ear/neck or chest) showed a greater probability of replying ‘Yes’ to heartbeat simultaneity judgments compared to those sampling from the left side of the chest. Participants’ confidence in their decision about simultaneity between heartbeat sensation and auditory tone presentation was maximal for the 0 and was lower after 100ms the ECG R-wave. Interestingly, higher confidence was related to reduced dispersion of sampling locations. We found evidence supporting the absence of relationship between temporal and spatial heartbeat sensations perception. Finally, we found evidence toward a relationship between spatial precision of heartbeat sensations and state anxiety score, which seems independent from the cardiovascular parameters.

## Discussion

Altogether, we find that, on average, heartbeat sensations occurred maximally 250ms after the ECG R-wave and were more frequently sampled from the left part of the chest. These findings, from our rigorous application of a multi-interval task, extend evidence from previous studies concerning interoceptive processing and conscious access to bodily signals (Brener et al. 1993, 1994; Khalsa et al. 2018; S. S. Khalsa et al. 2009; Ring and Brener 1992). Moreover, we observed that participants’ confidence in their experience of simultaneity judgement -between tones and their heartbeat sensations-was minimal for the 0 ms intervals and was lower after 100ms the ECG R-wave. Even though the left part of the chest was the most frequent location of heartbeat sensation, those individuals who felt heartbeats on the right of their upper body (head/ear/neck or chest) showed a greater probability of replying ‘yes’ to heartbeat simultaneity judgments compared to those sampling from the left side of the chest. Speculatively, the observed pattern of perceptual lateralization may have a basis in peripheral (left versus right vagus nerve) anatomy (Craig 2002) and central neural organisation where interoceptive inputs, integrated within right and the left anterior insula respectively, are putatively re-represented in the dominant right anterior insula (Craig 2002). Arguably, this may also mean that, in general, most participants base their judgments of cardiac timing and synchrony on spinothalamocortical information rather than vagus nerve afferents. However, such right-side dominance merits further evaluation, not least because it was not present for confidence ratings; higher confidence was observed for heartbeat sensations felt in the left head/ear/neck compared to left chest. Nevertheless, there remains some coherence with the hypothesis of right cerebral hemisphere engagement in the representation of heartbeat sensations attributable to peripheral cardiovascular asymmetries (Critchley et al. 2004; Katkin, Cestaro, and Weitkunat 1991; Katkin and Reed 1988). Confidence in heartbeat sensations may also be affected by pre-existing beliefs and biases arising from the participants’ understanding of anatomy (*e.g.* the heart is placed in the left part of the chest) and by quasi-interoceptive somatosensory pathways from the chest wall or the skin, which may afford great perceptual precision relative to viscerosensation. We have some evidence to support this second explanation, notably that higher confidence was associated with reduced dispersion of the sampling location; and, therefore, better spatial precision that suggests a potential somatosensory contribution. Indeed, it is accepted that interoceptive sensations such as visceral pain are poorly localized and may be felt at sites distal to than the affected organ due to overlap, recruitment and misinterpretation between somatic and visceral afferent information through shared relays (Cervero and Tattersall 1986; Giamberardino, Affaitati, and Costantini 2010; Sikandar and Dickenson 2012). Examples include shoulder pain associated with diaphragmatic involvement and left arm ache from cardiac angina.

A second question that we aimed to address was whether temporal and spatial perceptions of heartbeat sensation relate to each other. Simplistically one might predict that a greater sampling distance of heartbeat/pulse sensation from the heart would be associated with a greater lag in the perception of its timing, reflecting the inherent delay in the blood pulse wave’s activation of somatic mechanoreceptors and subsequent signaling to somatosensory/interoceptive cortices. In our present study, we found evidence against a simple relationship between timing of heartbeat sensations with both the distance from the heart and the dispersion of sampling location. In fact, our findings suggest the absence of any clear consistent relationship between the temporal and spatial perception of heartbeat sensations across individuals.

Our study further enabled us to test for the contribution of individual characteristics to heartbeat sensations. Physiologically, we showed that a slower heart rate was associated with a smaller preferred interval, but did not predict a specific timing of the actual heartbeat perception within the cardiac cycle (e.g. mean or median of intervals considered as in synch). This is interesting as both heart rate and HRV have previously been shown to influence performance in a multi-interval task, an effect proposes to arise because either people with slow heart rates have additional time to process cardiac sensations, or show differences in expectancies (Knapp-Kline and Kline 2005). Given the absence of a relationship between heart rate and the temporal precision of heartbeat perception, our results rather support the notion that heart rate expectancies exert a potentially greater impact on performance. Another important, and to a degree unexpected, finding within this study of non-clinical individuals was the absence of a relationship between trait anxiety and the timing of heartbeat sensations. Indeed, disrupted interoceptive ability is widely described in people suffering from anxiety disorders (Domschke et al. 2010; Garfinkel and Critchley 2013; Khalsa et al. 2018; Pang et al. 2019; Quadt et al. 2018) relationships between interoceptive accuracy and anxiety score are frequently described in ‘analogue’ populations (e.g. Dunn et al., 2010). Nevertheless, our results suggest that specific cardiac-timing paradigms can be implemented effectively in sub-clinical anxious populations since non-clinical anxiety does not seem to modulate the temporal perception of heartbeat sensations. However, we observed that higher levels of state anxiety were associated with lower spatial precision (increased sampling locations dispersion) even after controlling for cardiovascular parameters. Since people who feel their heartbeats reliably in the same anatomical location most likely are drawing upon somatosensory feedback (e.g. from the skin) rather than the less precise interoceptive feedback from viscerosensory afferents (Sahib S. Khalsa et al. 2009), our data suggest that state anxiety symptoms do not depend greatly on this somatosensory contribution to interoceptive experience.

Our results of this study should be considered in light of several constraints. First, headphones were used to deliver sounds to participants. Both ears are shown to be key areas of heartbeat sensation; areas not highlighted in limited previous research (Hassanpour et al. 2016; Khalsa et al. 2018; Sahib S. Khalsa et al. 2009; S. S. Khalsa et al. 2009). This may suggest that the pressure of the headphones may have given somatosensory feedback and influenced the location of heartbeat sensation experienced by participants. Also, as shown by our use of Bayesian statistics (Bayes factors), further studies in larger participant samples are required to test the relationship between interindividual characteristics and heartbeat sensations to generate firm conclusions. Nevertheless, our systematic investigation of the temporal and spatial perception of heartbeat sensations provides important fresh insights for the fields of experimental psychology, psychiatry and neuroscience. Further studies involving neuroimaging, ideally disentangling the contribution from both interoceptive and somatosensory pathways, will be helpful to build on this mechanistic understanding of embodiment, individual differences and the contribution of interoceptive signaling to emotion, cognition and behaviour.

## Methods

### Participants

Sixty-two volunteers (29 males, 33 females) aged from 18 to 45 years (*M* = 23.48 years, *SD* = 4.69) were recruited via advertisements at the University of Sussex and Brighton and Sussex Medical School. Given a medium effect size (eta partial square = 0.27 se Ring and Brener 1992), an alpha of 0.05, a beta of 0.85, a minimum of 48 participants needed to be recruited. We recruited more than 48 participants in anticipation of potential outliers.

All participants were healthy individuals with no history of psychiatric or neurological diseases and were not taking medication. One participant did not meet the inclusion Criteria and was excluded from the study. Participants were informed that they would complete a series of psychometric questionnaires and would take part in two tasks for one and a half hour. All participants gave their written informed consent and were compensated for their time (£15). The study was reviewed and approved by the BSMS Research Governance and Ethics committee.

### Demographic and psychometric description of the sample

The final sample was composed of 52 participants (26 Females, age: *M* = 22, *Median* = 22, *SD* = 4.7, *Range* = 18 - 45, years of education: *M* = 16.41, *Median* = 16, *SD* = 2.4, *Range* = 11-21). Characteristics of the sample and psychometric measures are presented in Table 4.

**Table 4.** Demographic and Psychometric measures: Minimum, 1^st^ quartile, Median, Mean, standard deviation, 3^rd^ quartile and maximum of Body Mass Index (BMI) and psychometric measures

|  | **BMI** | **BDI** | **STAI1** | **STAI2** | **TAS** |
| --- | --- | --- | --- | --- | --- |
| **Min** | 16.61 | 0.00 | 20.00 | 20.00 | 26.00 |
| **1st Qu.** | 20.31 | 2.00 | 25.75 | 32.00 | 41.75 |
| **Median** | 22.15 | 4.50 | 32.00 | 38.00 | 47.50 |
| **Mean** | 22.21 | 7.94 | 34.25 | 41.44 | 48.56 |
| **Sd** | 2.70 | 9.34 | 10.84 | 11.77 | 11.20 |
| **3rd Qu** | 23.73 | 9.50 | 41.25 | 50.25 | 56.00 |
| **Max.** | 30.78 | 43.00 | 64.00 | 65.00 | 72.00 |

### Procedure

The study was conducted in dedicated human testing facilities at the University of Sussex. Participants gave demographic information (e.g. age, weight, height) and a set of completed questionnaires, before the experiment. They next performed an audio-visual simultaneity task (for familiarisation with task demands) followed by the multi-interval heartbeat discrimination task that shared the same design structure.

### Questionnaires

#### *Beck Depression Inventory* (BDI)

The presence of depressive symptoms in participants was quantified using the Beck Depression Inventory (BDI). This questionnaire consists of 21 questions measuring cognitive, affective and somatic symptoms of depression as experienced by participants in the last 2 weeks. Each question is scored 0-3 with a higher number indicating a greater degree of symptom severity; allowing for a total score of up to 63. A score of 14-19 suggests mild depression, 20-28 suggests moderate depression and 29-63 indicates severe depression (Beck et al. 1996).

#### *Toronto Alexithymia Scale-20 items* (TAS-20)

The TAS-20 consists of 20 items rated on a five-point Likert scale (from 1 ‘strongly disagree’ to 5 ‘strongly agree’). The TAS-20 is composed of three factors (F1, F2, F3). The first-factor measures difficulties in identifying feelings (DIF), the second factor measures difficulties in describing feelings (DDF) and the third-factor measures the way the participant uses externally oriented thoughts (EOF). The total alexithymia score is the sum of responses across all 20 items. We considered the total score only in our analyses (Bagby, Parker, and Taylor 1994).

#### *Trait Anxiety Inventory* (STAI)

Trait anxiety was assessed using the Trait version of the Spielberger State/Trait Anxiety Inventory (STAI). This questionnaire is composed of 20 questions, assessing trait anxiety with questions such as “I lack self-confidence” and “I have disturbing thoughts”. Participants were asked to answer each statement using a response scale which runs from ‘Almost never’ to ‘Almost always’ to establish if there was a stable dispositional tendency (trait) for anxiety (Spielberger et al. 1983).

### Apparatus & task

Participants viewed a 24-inch monitor at an approximate distance of 50 cm. The monitor’s visual display had a screen resolution of 1920 × 1200 pixels and a refresh rate of 60 Hz. Auditory stimuli were delivered to participants through headphones. Experimental task procedures were implemented as in-house programmes using Psychophysics Toolbox Version 3 (http://psychtoolbox.org/) running in Matlab R2013a (The MathWorks, Inc., Natick, MA).

During the Multi-interval heartbeat discrimination task, tones were synchronised to specific time points within the cardiac cycle using electrocardiography (ECG) implemented via Cambridge Electronic Design (CED) hardware and Spike2 physiological recording software (version 7.18). Cardiac events were interfaced with the task events in Matlab. Three Ag/AgCl electrodes (3 M Healthcare, Neuss, Germany) were attached with foam tape: two on the upper left and right chest, and a ground electrode above the left hipbone. The ECG signal was sampled at 1000 Hz, amplified (1902, CED) and relayed to Spike2 recording software via an analogue-to-digital recorder (1401, CED). An inter-active threshold in the Spike2 recording isolated each ECG R-wave peak, which then primed tones delivery in the Matlab task script. The computer was equipped with a Strix Soar (https://www.asus.com/us/) soundcard allowing an input-output latency < 10ms and SNr> 110dB.

In the multi-interval heartbeat discrimination task, each participant was required to judge the simultaneity of a sequence of 5 tones with his/her own heartbeat (see Figure 1) (Brener et al. 1993; Brener and Ring 2016). This computerised task examines the participant’s ability to integrate interoceptive (heartbeat sensation) and exteroceptive (auditory stimuli) signals. Before the beginning of the task, the participant was instructed not to palpate his/her own pulse at any point during the experiment. Sequences of 5 tones were played to the participant, primed by his/her own ECG R-peaks. The tones of the sequence were delayed by one of the six time intervals (SOA: 0, 100, 200, 300, 400, 500ms) after the ECG R-peak. The participant had to decide whether the tones were played simultaneously with his/her own heartbeat or not and, then, rate how confident was that decision, using a visual analogue scale (VAS) on a computer screen. The participant was then presented with an outline image a body on the screen and had to click on the body where the heartbeat sensation was felt the strongest. Coordinates (x,y) of the selected body part were recorded. Each tone was played for a duration of 0.5s at a frequency of 48000Hz. The time interval between each question was equal to 1s and inter-trial interval (i.e. period between the trials) was equal to 5s. Overall, 120 trials were presented, with 20 trials per interval. These trials were completed over four separate 30-trial blocks with opportunities for rest in between blocks if required. The task lasted approximately 45 minutes. For two participants, we failed to record the location of heartbeat sensations. The familiarisation task (an audio-visual simultaneity task with the same design structure) is described in the supplementary section.

### Data analyses

Data were checked for outliers. One participant did not perform the audio-visual simultaneity task correctly (i.e. replied yes for all trials), four participants were not able to perform multi-interval heartbeat discrimination task (i.e. probability did not reach 0.50 of ‘yes’ response for any SOAs). These participants were removed from subsequent analyses. After computing the mean confidence and just-noticeable difference (described in the Supplementary section) for the audio-visual simultaneity task, participants whose performance data fell outside 1.5 times the interquartile range above the upper quartile and below the lower quartile (see whiskers’ boxplot) were labelled as outliers and their data was excluded (Chambers et al. 1983). The final sample size was equal to N=52 (including 2 participants without localisation data).

Body Mass Index (BMI; weight (kg) / height (m)^2^) was computed for each participant. All demographic, psychometric, and performance data for both tasks were held in long format (for mixed-effects effects models analyses) and short format in an averaged form (for correlations and descriptive statistics).

For each participant, the average interval (of reported heartbeat synchrony), its standard deviation, median and mode were calculated. The mode was used to assess the preferred time interval as being “in sync” with the heartbeat (i.e. temporal location of heartbeat sensation). The mean confidence and mean inter-beat interval (IBI) duration during the task were also computed. The standard deviation of the inter-beat interval (IBI SD) duration during the task was calculated and used as a marker of heart rate variability (HRV).

Concerning spatial perception of heartbeat sensation, for each trial, the distance between the sampling location and the heart (assigned a standardised location) was computed using coordinates marked by the participant on the body outline. Next, the mean of the distance to the heart was computed for each participant (Distance from the heart). We also computed dispersion from sampling locations by computing the mean of the standard deviation of X coordinates and the standard deviation of Y coordinates, for each participant ((sd(X) + sd(Y))/2). Finally, clusters of sampling location data points were defined using expectation-maximization (EM) algorithm for fitting mixture-of-Gaussian models (mclust R package) (Scrucca et al. 2016) and attributed to body parts and assigned names based on visual inspection.

### Statistical Analyses

All analyses were conducted in the R environment, version 3.6.1 (RCoreTeam 2013). Descriptive statistics were computed for all variables.

A non significant *p*-value is not enough to provide evidence toward the null hypothesis or toward the fact that the data are insensitive and that additional data are needed to conclude (Dienes 2014; Quintana and Williams 2018). Therefore, to facilitate the interpretation of our data, we ran separate Bayesian analyses, computing the Bayes Factor (BF) to indicate strength of evidence; P-values were used as the basis of decision making in respect of the compared hypotheses. Differences were considered significant when the probability *p* of a type I error was below 0.05.

Linear mixed-effects models were used in the analysis of confidence measures as the outcome was continuous. Generalized linear mixed models were used to analyse simultaneity assessement probability as the outcome was binary (non-simultaneous =0; simultaneous =1; binomial family function). In all models, participants were treated as a random factor with random intercepts (Barr et al. 2013). For frequentist analyses, the lme4 package was used (Bates et al. 2015) and p-values were computed using lmerTest package (Kuznetsova, Brockhoff, and Christensen 2014). 95% confidence intervals were computed and presented in each table. Two-sided contrasts were computed using the emmeans package ((Lenth et al. 2019) and p-values were corrected following the Holm–Bonferroni method. Based on previous work, we expected that participants would judge tones delivered between 100-300ms after initiation of ventricular contraction as simultaneous with their heartbeat (J. Brener et al., 1993; J. Brener & Kluvitse, 1988; Jasper Brener et al., 1994; C. Ring & Brener, 1992). Planned contrasts between delays from 100 to 300 ms were thus computed. We expected confidence to be related to difficulty and therefore to be maximal for the 0 and 500 intervals and minimal for the intervals between 100 to 300. Planned contrasts were computed.

Bayesian models were created in Stan computational framework (http://mc-stan.org/) accessed with the brms package (Buerkner 2017). To improve convergence and guard against overfitting, we specified weakly informative conservative priors (normal(0, 10)). Iterations were set to 2000 and chains to 4, where iteration numbers could be increased to achieve convergence. For each model and two-sided contrasts, Bayes Factor (BF) against the null, based on prior and posterior samples of a single parameter was estimated using the bayestestR package (Makowski, Ben-Shachar, and Lüdecke 2019). For contrasts, we also computed the most credible value and the 95% credible intervals (95% CrI in brackets), using brms package.

Two-sided frequentist Pearson correlations and partial correlations coefficients were calculated using Hmisc package (Harrel 2015) and Bayesian correlations, using BayesFactor package (https://richarddmorey.github.io/BayesFactor/).

A BF greater than 3 can be considered as substantial evidence against the null model, while a BF smaller than 1/3 indicates substantial evidence in favour of the null model (Wetzels et al. 2011).

Our interpretations required coherence between *p*-values and *BF*s (e.g. evidence for an effect was characterized by *p*-value < .05 and *BF* < 3).

### Data Availability Statement

The dataset and MATLAB scripts used in this study will be made available upon request by the corresponding author, Dr Sophie Betka.

## Supporting information

Supplementary material and results

## Acknowledgements

We thank Dr David Watson for his help with the set-up, as well as Dr Fosco Bernasconi and Dr Michael Pereira for their statistical advice. The project has been funded by the Rotary Foundation (Rotary Global Grant GG1526999).

## Author Contributions

Conceptualization (BS, ŁM, GS, CH), Methodology (BS, ŁM, GS, CH), Formal analysis (BS, CH), Software (BS, ŁM), Writing – original draft preparation (SB, CH), Writing – review and editing (BS, ŁM, GS, CH), Supervision (BS, ŁM,GS, CH), Project administration (BS, SM, KJ), Funding acquisition (GS, CH).

## Additional information & declaration of interests

The authors declare no competing interests.

