## Supplementary material and results for "Feeling the beat: Temporal and spatial perception of heartbeat sensations": Supp_MIT_v2.docx

### Supplementary results

### When do people perceive their heartbeats in relation to an external auditory tone and how confident are they?

|  | Average | Standard Deviation | Median | Mode | Confidence | IBI (s) | Distance from the heart | Sampling location dispersion |
| --- | --- | --- | --- | --- | --- | --- | --- | --- |
| Min | 196.00 | 122.30 | 200.00 | 0.00 | 3.53 | 0.59 | 20.22 | 4.35 |
| 1st Qu. | 241.60 | 161.80 | 200.00 | 100.00 | 48.22 | 0.74 | 59.92 | 16.79 |
| Median | 254.00 | 166.60 | 300.00 | 300.00 | 62.59 | 0.81 | 91.70 | 31.64 |
| Mean | 257.40 | 163.30 | 258.70 | 265.40 | 58.78 | 0.83 | 111.52 | 42.69 |
| Std | 31.31 | 13.56 | 55.77 | 149.36 | 19.32 | 0.12 | 71.93 | 36.95 |
| 3rd Qu | 268.70 | 171.70 | 300.00 | 400.0 | 72.11 | 0.89 | 150.96 | 54.33 |
| Max. | 353.70 | 180.00 | 400.00 | 500.0 | 97.60 | 1.06 | 373.63 | 153.72 |

**Table S1 Temporal perception of heartbeat sensations**: Minimum, 1^st^ quartile, Median, Mean, standard deviation, 3^rd^ quartile and maximum of average, standard deviation and Median of chosen delays (SOAs), as well as of confidence, inter-beat interval (in s), localisation distance from the heart and localisation dispersion.

| SOAs | β (Log odds) | *SE* | *z* value | *p*-value | 2.5% | 97.5% | *BF* |
| --- | --- | --- | --- | --- | --- | --- | --- |
| 0 (Intercept) | -0.01 | 0.14 | -0.05 | .961 | -0.20 | 0.20 | 0.01 |
| 100 | 0.33 | 0.10 | 3.32 | .001 | 0.14 | 0.49 | 0.51 |
| 200 | 0.59 | 0.11 | 5.27 | < .001 | 0.40 | 0.76 | 252.44 |
| 300 | 0.49 | 0.13 | 3.82 | < .001 | 0.31 | 0.67 | 76.25 |
| 400 | 0.46 | 0.15 | 3.07 | .002 | 0.28 | 0.64 | 69.53 |
| 500 | 0.29 | 0.18 | 1.63 | .104 | 0.11 | 0.46 | 0.29 |

**Table S2 Mixed-effects regression model to predict the effect of delays (SOAs) on probability to say “yes”**, with random intercepts, including all participants. For each level of the parameter, log odd, standard error (*SE*), *z*-value, *p*-value, 95% confidence interval and Savage-Dickey density ratio Bayes Factor (*BF*) are presented.

(Model_accuracy <- glmer(Jud ~ SOAsF + (1 | pxID), data = dat_long, family = binomial, control = glmerControl (optimizer="bobyqa", optCtrl = list(maxfun = 100000)));

Model_accuracy <-brm(Jud ~ SOAsF +(1|pxID),data =dat_long, family=bernoulli("logit"), prior = set_prior('normal(0,10)'), iter = 2000, chains=4).

N.B.: Log odds values (beta) can be converted back into proportions using the inverse logit formula Exp(beta)/(1+ exp(beta))

| SOAs | β | SE | df | t value | p-value | 2.50% | 97.50% | BF |
| --- | --- | --- | --- | --- | --- | --- | --- | --- |
| (Intercept) | 61.04 | 2.74 | 56.00 | 22.26 | < .001 | 55.62 | 66.46 | >1000 |
| SOAs 100 | -0.40 | 0.91 | 6169.00 | -0.44 | .658 | -2.18 | 1.38 | >1000 |
| SOAs 200 | -2.65 | 0.91 | 6169.00 | -2.91 | .004 | -4.43 | -0.87 | >1000 |
| SOAs 300 | -3.65 | 0.91 | 6169.00 | -4.01 | < .001 | -5.43 | -1.87 | >1000 |
| SOAs 400 | -3.10 | 0.91 | 6169.00 | -3.41 | .001 | -4.88 | -1.32 | >1000 |
| SOAs 500 | -3.75 | 0.91 | 6169.00 | -4.12 | < .001 | -5.53 | -1.97 | >1000 |

**Table S3 Mixed-effects regression model predicting the effect of delays (SOAs) on confidence ratings**, with random intercepts, including all participants. For each level of the parameter, log odd, standard error (*SE*), degree of freedom (*df*), *t*-value, *p*-value, 95% confidence interval and Savage-Dickey density ratio Bayes Factor (*BF*) are presented.

(Model Confidence <- lmer(Conf ~ SOAsF + (1 |pxID), data = dat_long, control = lmerControl(optimizer="bobyqa", optCtrl = list(maxfun = 100000)));

Model Confidence <- brm(Conf ~ SOAsF + (1 |pxID), data = dat_long, family=gaussian , prior = set_prior('normal(0, 10)', class = 'b'), sample_prior = TRUE,iter = 10000, chains=4, save_all_pars = TRUE)).

**Where do people feel their heartbeat and how confident are they?**

| Cluster | β | SE | z.value | *p*.value | 2.50% | 97.50% | BF |
| --- | --- | --- | --- | --- | --- | --- | --- |
| Left part of the chest (Intercept) | 0.26 | 0.10 | 2.59 | 0.010 | 0.06 | 0.46 | 0.26 |
| Left part of the head/ear/neck | -0.12 | 0.12 | -0.98 | 0.328 | -0.35 | 0.12 | 0.02 |
| Right part of the head/ear/neck | 0.37 | 0.11 | 3.29 | 0.001 | 0.15 | 0.59 | 720.06 |
| Right part of the chest | 0.48 | 0.14 | 3.38 | 0.001 | 0.21 | 0.77 | 273.05 |
| Left fingers | 0.25 | 0.18 | 1.40 | 0.163 | -0.10 | 0.61 | 0.43 |
| Miscellaneous | -0.03 | 0.17 | -0.16 | 0.874 | -0.36 | 0.31 | 0.03 |
| Right fingers | -0.47 | 0.27 | -1.65 | 0.098 | -1.04 | 0.08 | 0.02 |
| Left arm | -0.31 | 0.39 | -0.79 | 0.431 | -1.09 | 0.46 | 0.03 |

**Table S4 Mixed-effects regression model predicting the effect of spatial clusters on simultaneity judgements,** with random intercepts, including participants with localisation data (N=50). For each level of the parameter, log odd, standard error (*SE*), *z*-value, *p*-value, 95% confidence interval and Savage-Dickey density ratio Bayes Factor (*BF*) are presented.

(Model Accuracy Cluster <- glmer(Jud~ cluster_newF + (1 |pxID), dat_long2, family= binomial, control = glmerControl(optimizer="bobyqa", optCtrl = list(maxfun = 100000)));

Model Accuracy Cluster <- brm(Jud~ cluster_newF + (1 |pxID), data = dat_long2, family=bernoulli("logit"), prior = set_prior('normal(0, 10)', class = 'b'), sample_prior = TRUE,iter = 2000, chains=4, save_all_pars = TRUE).

N.B.: Log odds values (beta) can be converted back into proportions using the inverse logit formula Exp(beta)/(1+ exp(beta))

| Cluster | β | SE | df | t.value | *p*.value | 2.50% | 97.50% | BF |
| --- | --- | --- | --- | --- | --- | --- | --- | --- |
| Left part of the chest (Intercept) | 55.75 | 2.52 | 54.32 | 22.10 | <.001 | 50.78 | 60.73 | >1000 |
| Left part of the head/ear/neck | 8.155 | 1.30 | 5709.29 | 6.25 | <.001 | 5.6 | 10.70 | >1000 |
| Right part of the head/ear/neck | 5.70 | 1.17 | 5825.64 | 4.89 | <.001 | 3.41 | 7.98 | >1000 |
| Right part of the chest | 6.33 | 1.51 | 5715.00 | 4.21 | <.001 | 3.39 | 9.28 | >1000 |
| Left fingers | 3.42 | 1.88 | 5883.16 | 1.82 | .068 | -0.27 | 7.09 | >1000 |
| Miscellaneous | 1.14 | 1.77 | 5888.50 | 0.65 | .518 | -2.33 | 4.60 | >1000 |
| Right fingers | 5.32 | 3.18 | 5572.23 | 1.68 | .094 | -0.90 | 11.53 | >1000 |
| Left arm | 7.16 | 4.46 | 5363.83 | 1.60 | .109 | -1.57 | 15.92 | >1000 |

**Table S5 Mixed-effects regression model predicting the effect of spatial clusters on confidence ratings,** with random intercepts, including participants with localisation data (N=50). For each level of the parameter, log odd, standard error (SE), degree of freedom (df), t-value, p-value, 95% confidence interval and Savage-Dickey density ratio Bayes Factor (BF) are presented.

(Model Confidence Cluster <-lmer(Conf~ cluster_newF + (1 |pxID), dat_long2, family= binomial, control =lmerControl(optimizer="bobyqa", optCtrl = list(maxfun = 100000)));

Model Confidence Cluster <- brm(Jud~ cluster_newF + (1 |pxID), data = dat_long2, family=bernoulli("logit"), prior = set_prior('normal(0, 10)', class = 'b'), sample_prior = TRUE,iter = 2000, chains=4, save_all_pars = TRUE)

***Individual performances***

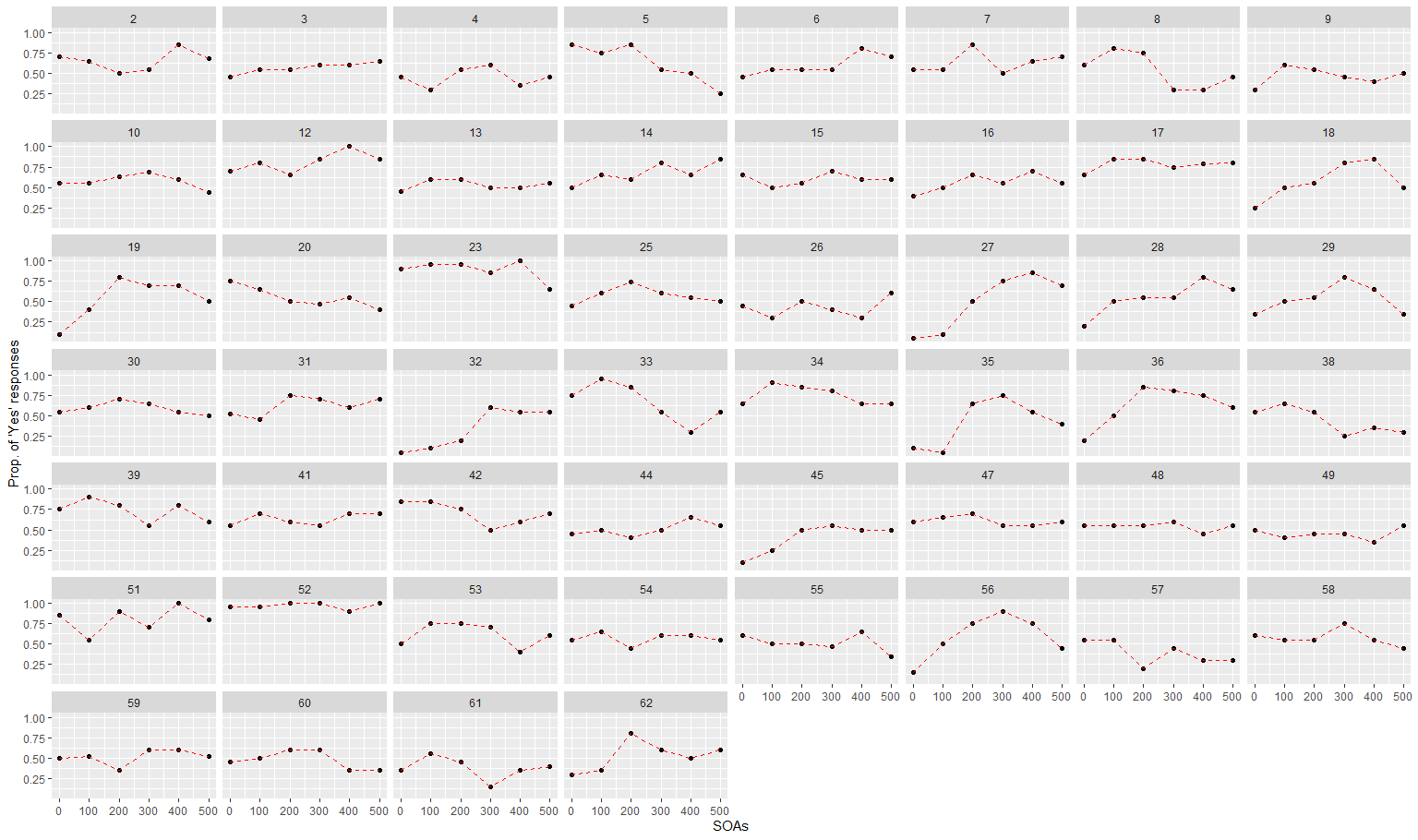

Figure S1 Temporal perception of heartbeat sensation for each participant.

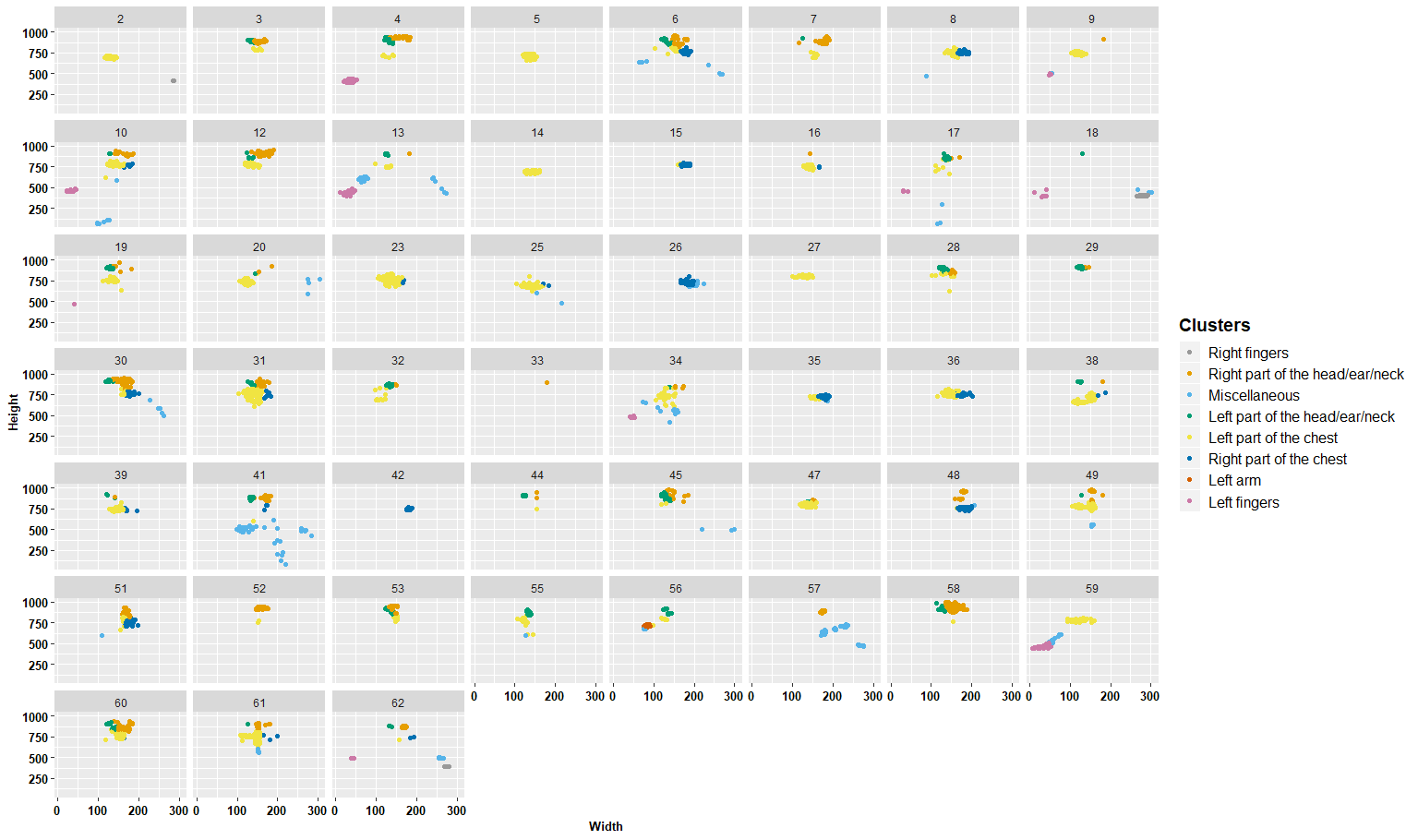

Figure S2 Clusters and sampling dispersion for each participant
